# Computational identification and in vitro evaluation of astragaloside II as an inhibitor of dengue virus NS1-induced endothelial hyperpermeability

**DOI:** 10.64898/2026.11.12.766166

**Authors:** Wanida Chawnawa, Natthawadee Leephatarakit, Warun Maneepitasut, Sarunya Maneerattanasak, Pongpak Manoret, Korawit Poyseungnoen, Chompunuch Boonarkart, Chunya Puttikhunt, Panisadee Avirutnan, Sansanee Noisakran, Prasert Auewarakul, Kobchai Duangrattanalert, Chanati Jantrachotechatchawan

**Affiliations:** Siriraj Center of Research Excellence in Dengue and Emerging Pathogens, Faculty of Medicine Siriraj Hospital, Mahidol University, Bangkok, Thailand; Molecular Biology of Dengue and Flaviviruses Research Team, Medical Molecular Biotechnology Research Group, National Center for Genetic Engineering and Biotechnology, National Science and Technology Development Agency, Bangkok, Thailand; Emerging Infectious Diseases Research Unit, Faculty of Medicine Siriraj Hospital, Mahidol University, Bangkok, Thailand; Faculty of Medicine Siriraj Hospital, Mahidol University, Bangkok, Thailand; AI-XENIX Co., Ltd., AI Solutions and Analytics, Bangkok, Thailand; Research Department, Faculty of Medicine Siriraj Hospital, Mahidol University, Bangkok, Thailand; Department of Microbiology, Faculty of Medicine Siriraj Hospital, Mahidol University, Bangkok, Thailand; Division of Dengue Hemorrhagic Fever Research, Research Department, Faculty of Medicine Siriraj Hospital, Mahidol University, Bangkok, Thailand; Department of Biology, Duke University, Durham, NC, USA

## Abstract

**Background:** Dengue is a mosquito-borne viral disease in which severe cases are marked by leakage of plasma from blood vessels. The secreted nonstructural protein 1 (NS1) of dengue virus damages the endothelial barrier and is a drug target, but no approved antiviral exists. We aimed to identify compounds that target NS1 and to test whether the selected compound, astragaloside II (ASII), counteracts NS1-induced endothelial hyperpermeability.

**Methodology/Principal findings:** We built a deep learning model that predicts compound– target binding from chemical fingerprints and protein sequences, and used it to prescreen 177 natural compounds against NS1 of the four dengue virus serotypes. The model outperformed the DeepConv-DTI baseline in recall and F1-score on a BindingDB holdout set and on two independent test sets. ASII was predicted to bind NS1 and was docked onto the NS1 dimer. In human microvascular endothelial cells (HMEC-1), NS1 reduced the relative trans-endothelial electrical resistance to 0.56 (P < 0.001), whereas ASII at 10–30 µg/mL kept resistance close to control levels without detectable cytotoxicity. ASII did not reduce NS1 internalization. In a competitive enzyme-linked immunosorbent assay, ASII inhibited NS1 binding to heparan sulfate in a dose-dependent manner (half-maximal inhibitory concentration 51.12 µg/mL; 95% confidence interval 47.91–56.24 µg/mL), although inhibition at the protective doses was only 8–26%.

**Conclusions/Significance:** ASII is a candidate for further study as a modulator of NS1-induced endothelial hyperpermeability. Its mechanism remains unresolved and requires further study. Direct binding to NS1, activity in infected cells and effects in other endothelial models were not tested here, and the computational predictions require experimental confirmation.

**Author summary:** Dengue is a viral disease spread by mosquitoes that infects hundreds of millions of people each year. In severe cases, fluid leaks out of the smallest blood vessels, which can cause shock and death. No approved medicine treats the infection itself. A viral protein called NS1, which infected cells release into the blood, helps cause this leakage by damaging the lining of the blood vessels. We therefore looked for a compound that could block the harmful effects of NS1. We trained a computer program to predict which chemicals are likely to stick to a given protein, and we used it to rank 177 natural compounds. We chose one of the best-ranked compounds, astragaloside II, which comes from a plant used in traditional Chinese medicine. In cultures of blood vessel cells, NS1 weakened the cell layer, and astragaloside II prevented this damage without harming the cells. The compound also reduced the binding of NS1 to a sugar-like molecule on cell surfaces, but only weakly at the doses that protected the cells. We did not test it in infected cells or animals, so we regard it as a lead for further work, not as a treatment.

## Introduction

Dengue virus (DENV), a mosquito-borne flavivirus, infects nearly 390 million people worldwide annually, leading to 500,000 hospitalizations and 20,000 deaths [1]. Infection with any of the four DENV serotypes (DENV-1, -2, -3, and -4) can cause a range of illnesses, from subclinical infection to dengue fever, and life-threatening dengue hemorrhagic fever and dengue shock syndrome. Severe dengue is associated with a transient increase in endothelial permeability, leading to vascular leakage [2]. Extensive leakage into tissue spaces and serous cavities, including the pleura, pericardium, and peritoneal cavities, along with hemorrhaging, can cause profound shock that may be fatal if not promptly managed [3]. Currently, no clinically approved antiviral treatment for dengue is available, and management relies on supportive therapy, primarily fluid management [1], as the underlying mechanisms of endothelial hyperpermeability remain unclear.

The DENV nonstructural protein 1 (NS1) is the viral protein released by infected cells and circulates at high levels in patients with severe dengue [4]. NS1 impairs the barrier function of endothelial cell monolayers by disrupting tight and adherent junctions [5] and degrading the endothelial glycocalyx layer (EGL) [6]. The EGL comprises glycoproteins with acidic oligosaccharides and terminal sialic acid, as well as proteoglycans with glycosaminoglycan side chains, including heparan sulfate and chondroitin sulfate [7]. NS1 binds to uninfected endothelial cells via these glycosaminoglycan [8], activating endothelial sialidases, heparanase, and cathepsin L [6]. This leads to glycocalyx degradation, which correlated with vascular leakage in severe DENV infection [9]. Additionally, NS1 is recognized by Toll-like receptor 4 (TLR4), activating macrophages and human peripheral blood mononuclear cells (PBMCs), which release proinflammatory cytokines that contribute to vascular leakage [10]. NS1 also directly activates endothelial cells via TLR4, further disrupting the monolayer and promoting vascular leakage [10].

Drug virtual screening by protein-ligand interaction modeling is of major importance for drug-discovery. The gold standard procedures are based on binding surface simulation including molecular docking which could require considerable amounts of time for a large number of candidates. Multiple deep learning models predict binding affinity from three-dimensional protein structure information e.g., atomic coordinates [11], spatial graph representation of covalent and non-covalent interactions [12], and 3D protein structure surface. However, these models still require computational resources for data processing and lack flexibility in terms of input data variation. One promising alternative is a model that uses protein primary sequence and chemical fingerprint representation such as DeepConvDTI [13], which accommodates manipulation of sequence inputs and allows for prediction on multiple variants of the same protein. Herein, we developed a model that applies a multiscale convolutional neural network (CNN) as does DeepConvDTI but in skip connection repeat to enhance learning of protein hierarchical structure and also incorporates multiple units of dropout and normalization to improve generalizability across protein sequences.

Given the critical role of NS1 in dengue pathogenesis, candidate compounds that target NS1 may reduce disease severity. A recent study has shown that NS1 inhibitors can effectively interfere with its function [14]. However, despite ongoing research, no antiviral drug has yet been approved for the treatment of dengue. This study aimed to identify compounds with inhibitory effects on DENV NS1 activity using both *in silico* and *in vitro* approaches.

## Methods

### Ethics statement

This study used an established human microvascular endothelial cell line and recombinant proteins. It did not involve human participants, human tissue or animals.

### Datasets

The main data of 822,260 protein-ligand pairs with IC_50_ as binding affinity for model training, validation, and evaluation was from BindingDB. Given that each protein-ligand pair may have multiple entries of IC_50_, any pairs with coefficient of variation (standard deviation/mean) of IC_50_ higher than 50% were excluded. The data was divided into 3 non-overlapping subsets – training, validation, and evaluation – of approximately 29:1:1 ratio such that the same protein or compound was not shared across different subsets. The manually curated dataset IUPHAR/BPS [15] Guide to Pharmacology, jointly developed by the International Union of Basic and Clinical Pharmacology (IUPHAR) and the British Pharmacological Society (BPS), and the dataset COVID Moonshot consortium in the BindingDB for non-covalent inhibitor were used as independent evaluation sets. The total number of pairs, unique proteins, and unique ligands, were listed in Table 1.

**Table 1.** Datasets for model training and evaluation.

| **Dataset** | **Subset** | ***N* pairs** | ***N* proteins** | ***N* ligands** |
|---|---|---|---|---|
| BindingDB | Train | 769,157 | 4,936 | 540,718 |
|  | Validation | 26,650 | 221 | 26,459 |
|  | Evaluation | 26,453 | 229 | 26,330 |
| IUPHAR/BPS | Independent | 222 | 123 | 195 |
| COVID Moonshot | Independent | 1,331 | 1 | 1,331 |

BindingDB subsets share no protein or ligand. IUPHAR/BPS and COVID Moonshot were used as independent evaluation sets.

### Data preprocessing

Protein sequences of length between 50 to 3,000 amino acids were tokenized using a vocabulary of 20 standard amino acids. Any proteins with non-canonical residues such selenocysteine or hydroxyproline were excluded. Ligand SMILES are transformed to their canonical SMILES form and subsequently converted to binary Morgan fingerprint of 1,024 bits at radius 2 using RDkit.

The hard threshold of IC_50_ differentiating the two labels of “Strong binding” and “Weak binding” was set at 100 nM or 10^-7^ M (pIC_50_ = 7). A standard sigmoid centered at the hard threshold was then applied to generate continuous soft labels from 0 to 1 as training outputs.

### Model architecture

The architecture of this model (Fig 1) was inspired by that of DeepConv-DTI. The protein sequence encoder is a stacked multi-scale convolutional neural network (CNN). It contains 3 sequential blocks of paralleled 1D convolutional layers of different kernel sizes (k = 10, 15, 20, 25, and 30) with add skip connections. Stacking enables extraction of hierarchical features. Multi-scale by different kernel sizes helps extract spatial features at varying scales. Spatial dropout randomly removes an entire feature map. This forces the network to avoid becoming overly reliant on any single feature detector and to learn more diverse and robust set of features to still converge at the correct prediction. A skip connection for each block prevents information loss or vanishing gradient. The small ligand molecular encoder consists of dense layers and Batch normalization and mish.

**Fig 1.**
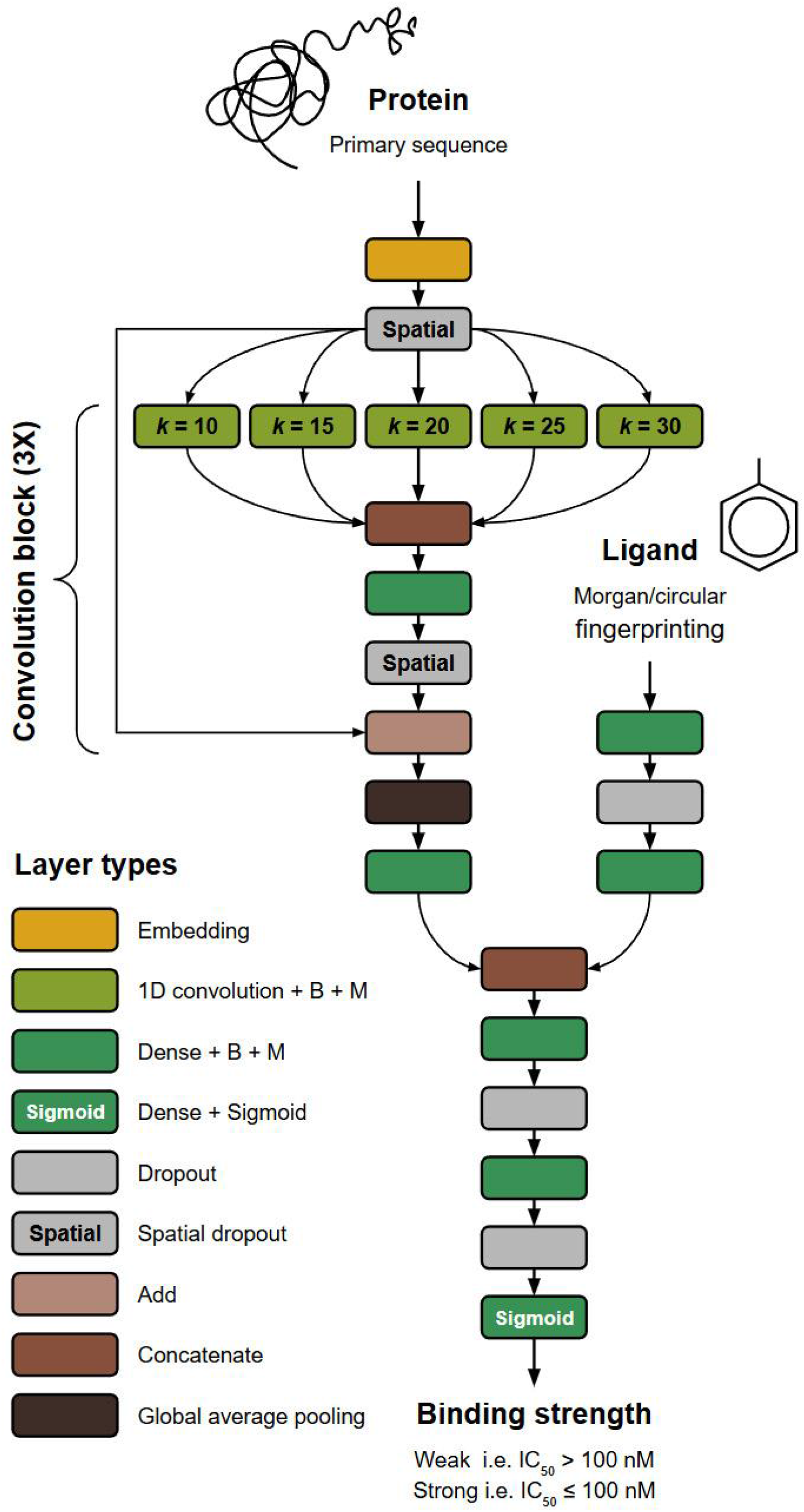
The architecture of the stacked multiscale CNN model. k, convolution kernel size; B, batch normalization; M, Mish activation.

### Training strategy

The current model and DeepConv-DTI, as a control, were trained on the same dataset. In each training step, input samples are stratified by labels and protein families. The models were trained for 500 epochs. The final weight sets were selected from the epoch of least total loss on the validation set.

### Post-modeling analysis

Performances of the models were evaluated on the main evaluation subset and the two independent datasets with the following metrics: precision, recall, F1-score, and area under curve of precision-recall (AUC-PR). Predictions can be one of the 4 types, actual strong binding correctly predicted (true positive: TP), actual weak binding correctly predicted (true negative: TN), weak binding mis-predicted as strong (false positive: FP), and strong binding predicted as weak (false negative: FN). Precision or positive predictive value (PPV) is TP/(TP + FP), whereas recall or sensitivity is TP/(TP + FN). F1-score is the harmonic mean between precision and recall on equal weights. AUC-PR is the area under the curve of precision-recall at different thresholds. Prediction scores were converted by a modified hyperbola with fixed intersections at (0,0) and (1,1) such that the initial classification threshold was transformed to 0.5.

To confirm the generalizability of the model, protein-ligand pairs of the holdout evaluation sets were divided into groups based on discretized range of protein lengths and sequence similarity with the proteins in the training set computed as Levenshtein distance. Correlation coefficients were then measured to indicate association between model performances in AUC-PR and group representative values of length or sequence similarity. Model performances were also calculated for protein-ligand pairs grouped by major protein families and the statistical differences were measured using one-way ANOVA.

### Molecule prescreening

A total of 177 compounds with valid canonical SMILES from a total of 477 natural compounds listed by Sigma-Aldrich were input to our model for their predicted binding against NS1 sequence of all four DENV serotypes.

### Molecular docking and structure prediction

CBdock2 [16] was used to explore the binding of the pre-screened candidate on the whole NS1 dimer structure (PDB: 4O6B [17]). Afterward, to verify this binding site identified by CBDock2, SwissDock Attractive Cavities 2.0 algorithm [18] was used to model surface docking of also the 3 controls C012 (PG545), C01, and C12. The number of the control compounds indicates specific moieties: 0 steroid backbone, 1 glucose tetramer backbone, and 2 sulfate groups on glucose tetramer. The 3 compounds were experimentally validated by pre-treatment with the compounds prior to NS1 treatment on primary human peripheral blood mononuclear cells (PBMCs) followed by IL-6 quantification by ELISA [19]. In the study, the IC50 values for C012 and C12 were approximately 211 nM and 29.2 μM respectively. On the other hand, C01 does not inhibit NS1-induced IL-6 release by PBMCs. The inhibition was verified to be caused by NS1 binding as none of the compounds elicited any blocking effects upon LPS-induced IL-6 release in the same set of experiment. Swissdock was used to validate the binding because sulfate group-containing C012 and C12 have the numbers of possible bond rotations that exceed the limits set by CBDock2. In turn, the required computational time to simulate binding of most molecules to at least cover half of the entire NS1 dimer and the regions of overlaps between the monomers also exceed the limits of Swissdock as well.

For the preliminary hypothesis test that the chosen candidate inhibits the binding between NS1 and the speculated target protein, sequences of DENV-2 NS1 and the target protein are input to the web application of AlphaFold3 [20] by Google which can predict the structure of the binding complex between the proteins. The copy number of NS1 sequence is set at 2 while the hypothetical target is at 1. Subsequently, the binding region of the strongest binding configuration predicted by CB-Dock2 is then highlighted on this complex to check for the superimposition between the ligand binding and the protein-protein binding.

The amino acid residues responsible for potential binding with heparan sulfates were identified by using Protein-Ligand Interaction Profiler (PLIP) web tool [21] on the 3D structure of NS1_172-352_ and soluble sulfates (PDB: 4OIG [22]).

### Cell culture

The human microvascular endothelial cell line-1 (HMEC-1: a gift from Professors Michael S. Diamond and John P. Atkinson, Washington University, St. Louis, USA) was maintained in MCDB 131 medium (Invitrogen) supplemented with 10 ng/mL human epidermal growth factor recombinant protein (Invitrogen), 1 µg/mL hydrocortisone (Sigma-Aldrich), 10% fetal bovine serum (Gibco), 1X GlutaMAX (Gibco), and 50 U/mL penicillin-streptomycin (Gibco). Cells were incubated at 37°C with 5% CO_2_.

### Trans-endothelial electrical resistance (TEER) assay

The effect of ASII on NS1-induced endothelial hyperpermeability was evaluated by measuring the TEER of HMEC-1 monolayers grown on a 24-well Transwell polycarbonate membrane system (Corning). Briefly, 5×10⁴ HMEC-1 cells were seeded into the apical side of the Transwell inserts (top chamber) in a final volume of 200 μL/well. Each Transwell was placed inside a 24-well plate containing 500 μL of complete MCDB 131 medium in the basolateral compartment and incubated at 37°C for 24 hours. Subsequently, NS1 from DENV-2 (10 μg/mL) with or without varying concentrations of ASII (10, 20, and 30 μg/mL; MedChemExpress, HY-N0433), was added to the apical side of the Transwell insert containing the cell monolayer. TEER values, expressed in Ohms (Ω), were measured at sequential time points of 0, 6, 9, and 24 hours after NS1 exposure using an epithelial volt-ohm meter and chopstick electrodes (Millicell^®^ ERS-2, Merck). NS1-exposed HMEC-1 treated with heparin (10, 20, and 30 μg/mL) served as positive controls for NS1-heparin binding. Control HMEC-1 cells that were neither exposed to NS1 nor treated with ASII, grown on Transwell inserts, were used as control cells, while inserts with medium alone served as blank resistance measurements. Endothelial permeability was expressed as relative TEER, with a value of 1.0 indicating normal permeability. The relative TEER was calculated as the ratio of resistance values using the following formula: [Ω experimental condition - Ω medium alone]/[ Ω control endothelial cells - Ω medium alone].

### Adenosine triphosphate (ATP)-based cell viability assay

The cytotoxic effect of ASII on HMEC-1 was examined by measuring intracellular ATP as an indicator of cell viability. Cells were seeded into 96-well plates at a density of 5×10⁴ cells/well in a fixed volume of 100 µL and incubated under 5% CO₂ at 37°C for 24 hours. Subsequently, the cells were treated with different concentrations of ASII (5, 10, 20, and 30 μg/mL), maintaining a final volume of 100 µL/well. After a 24-hour incubation, whole-cell lysates were used to evaluate ATP levels using the ATPlite Luminescence Assay System (PerkinElmer), following the manufacturer’s instructions. Briefly, 50 µL/well of mammalian cell lysis solution was added to inactivate endogenous ATPases, and the plate was shaken at 700 rpm for 5 minutes. Then, 50 µL/well of substrate solution was added and incubated for 10 minutes in the dark. The luminescent signal, measured using a microplate reader (Synergy H1, BioTek), was positively correlated with ATP levels. Cell viability was assessed as a percentage compared to control cells that were not treated with ASII.

### NS1 internalization assay

Since NS1 can be endocytosed by HMEC-1 *in vitro*, we investigated whether ASII reduces intracellular NS1 accumulation in these cells using an NS1 internalization assay. HMEC-1 cells were seeded into an 18-well chamber slide at 5×10⁴ cells/well in 100 µL of complete MCDB 131 medium and incubated at 37°C with 5% CO₂ for 24 hours. Following incubation, 100 µL of medium was discarded, and cells were treated with 100 µL of NS1 (10 or 20 μg/mL) with or without ASII (20 or 30 μg/mL) and incubated for 9 hours at 37°C. After incubation, 100 µL of supernatant was removed, and cells were washed once with 150 µL of complete MCDB 131 medium, followed by a second wash with 150 µL of diluent buffer (PBS supplemented with 2% FBS). Cells were then fixed with 100 µL of 4% paraformaldehyde in PBS for 20 minutes, washed with diluent buffer, permeabilized with 100 µL of 0.2% Triton X-100 in PBS for 15 minutes, and blocked with 100 µL of 5% bovine serum albumin (BSA) in PBS for 1 hour at room temperature. Then, cells were incubated with 50 µL of mouse anti-DENV NS1 mAb 1F11 for 1 hour and washed three times with diluent buffer. Subsequently, cells were incubated with 50 µL of Cyanine3-conjugated goat anti-mouse IgG (Jackson ImmunoResearch) and Hoechst (a specific nuclear stain) for 1 hour in the dark. After three additional washes with diluent buffer, 100 µL of 0.1% sodium azide in PBS was added to each well. Finally, stained cells were analyzed using a confocal microscope (Zeiss). The levels of NS1 inside the cells were quantified by analyzing at least 25 different fields with Zen software (Zeiss) and expressed as the sum of fluorescence intensity from each field.

### Competitive binding assay

A competitive enzyme-linked immunosorbent assay (ELISA) was performed to assess the effect of ASII on the interaction between NS1 and heparan sulfate (HS), as NS1 is known to bind HS on the surface of endothelial cells [8]. A microplate (Thermo Fisher Scientific) was coated with 50 µL/well of 1 mg/mL HS (MedChemExpress, HY-101916) in PBS and incubated overnight at 4°C, as previously described [8]. After four washes with PBS, unbound sites were blocked with 100 µL/well of 2% BSA (Sigma-Aldrich) in PBS at 37°C for 2 hours. Following five washes with PBS containing 0.05% Tween-20 (PBST), 3 μg/mL DENV-2 NS1 was mixed with an equal volume of varying concentrations of ASII (1-100 μg/mL) or 2.4% DMSO, added to the wells, and incubated at 37°C for 2 hours. The plate was then washed five times with PBST, followed by a 1-hour incubation at 37°C with 50 μL/well of anti-DENV NS1 monoclonal antibody (2G6). After washing, horseradish peroxidase-conjugated goat anti-human IgG antibody (Merck) was added for a 1-hour incubation at 37°C. Following six washes with PBST, the signal was detected by adding 50 μL of TMB substrate (SeraCare), followed by 50 μL of 1 M H_2_SO_4_ stop solution to each well. Absorbance was measured at 450 nm using a microplate reader (Synergy H1, BioTek). Assays for each condition were performed in duplicate across two independent experiments. Absorbance values were converted to percent inhibition using the formula: (1 − B/B_0_) × 100, where B is the absorbance value of a tested sample solution, and B_0_ is the absorbance value of a similar solution without ASII.

### Statistical analysis

All statistical analyses and graphs were generated using GraphPad Prism version 9.0 (GraphPad Software Inc., California, USA). Data are presented as the mean ± standard error of the mean (SEM). An unpaired t-test was used to compare two independent groups. For TEER experiments, an ordinary two-way analysis of variance (ANOVA) was performed to assess differences among all experimental and control groups, followed by Dunnett’s test for post-hoc comparisons between the experimental groups and the control group. The half-maximal inhibitory concentration (IC_50_) value was determined from a dose–response curve using nonlinear regression. A *P*-value < 0.05 was considered statistically significant.

## Results

### Model performance across datasets

The model was trained, validated and evaluated on protein–ligand pairs from BindingDB [23] and was tested on two independent sets, IUPHAR/BPS Guide to Pharmacology [15] and the COVID Moonshot consortium set. The numbers of pairs, unique proteins and unique ligands are given in Table 1.

The architecture of this model (Fig 1) was inspired by that of DeepConv-DTI [13] except that instead of having one multi-scale convolutional neural network (CNN) unit, the protein encoder of this model contains a stack of 3 sequential blocks of paralleled 1D convolutional layers of different kernel sizes with additive skip connections. Stacking enables extraction of hierarchical features of the protein sequence spatial features of varying scales that have already been captured by multi-scale CNN with different kernel sizes. Furthermore, this model also adopts multiple techniques and layers to enhance the overall robustness and reduce overfitting. Spatial dropout, by randomly removing an entire feature map, forces the network to avoid becoming overly reliant on any single feature detector and to learn more diverse and robust set of features to still converge at the correct prediction. A skip connection for each multi-scale CNN stack prevents information loss or vanishing gradient. Mish activation function, unlike standard ReLU, is continuously differentiable, encouraging stable gradients and weight updates during backpropagation [24]. Mish is non-monotonic with negative value for x < 0 and also slower convergence to the y = 0 asymptote. This allows for a control of model complexity and regularization.

To account for potential class imbalance from a smaller number of positive pairs, focal cross-entropy loss was used to give relatively higher priority to the difficult class. For weight updates, rectified Adam (RAdam) was used as an optimizer [25]. The standard Adam optimizer enables different degrees of weight updates across layers, providing flexibility and often leading to faster optimization. However, the variance in weight updates across layers can become too large and cause instability in model training. To address this, RAdam optimizer was developed to stabilize training by reducing the variance of weight updates across layers.

For the holdout BindingDB evaluation set, our current model performed better than the reference DeepConv-DTI especially in recall with negligible decrease in precision (Fig 2), resulting in greater overall performances of both AUC-PR and F1-score. For the IUPHAR independent set, our model also outperformed in recall, AUC-PR, and F1. For the COVID MC set, our model outperformed DeepConv-DTI in all metrics.

**Fig 2.**
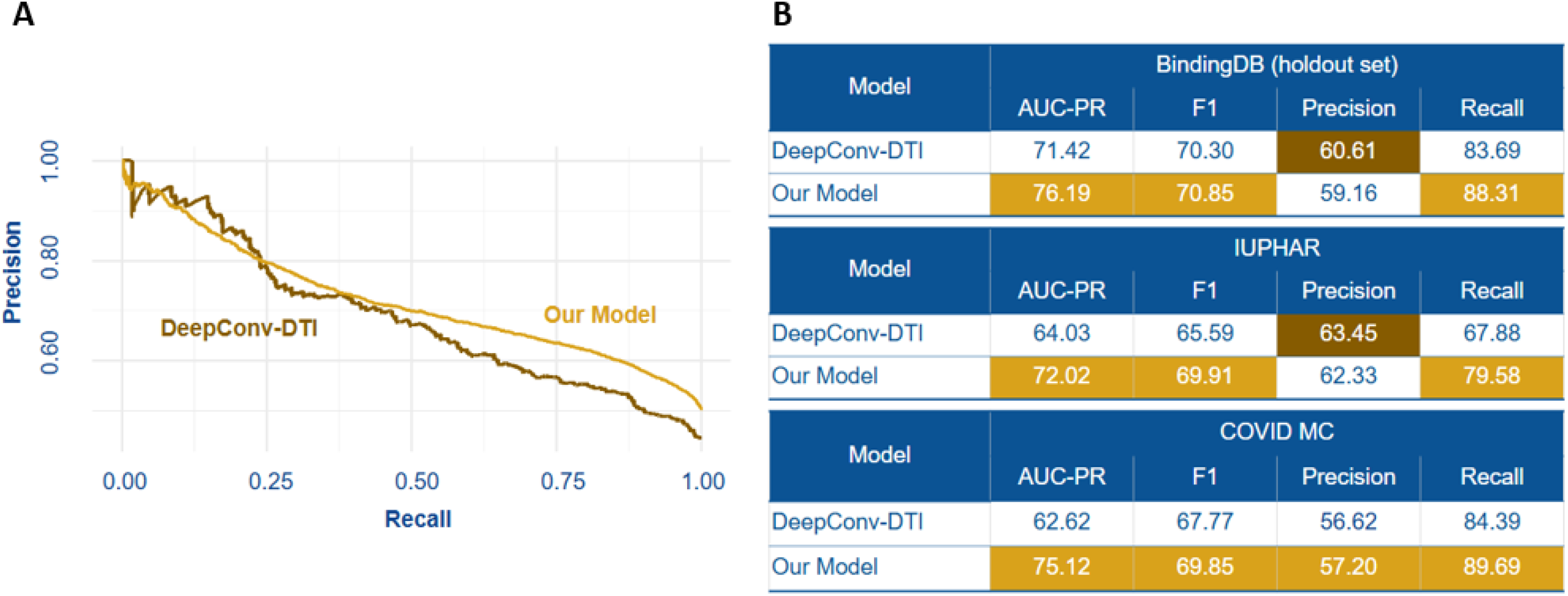
Comparison between the performances of our model and DeepConv-DTI. (A) Precision-Recall curves between the two models on BindingDB holdout set and (B) Classification report with AUC-PR, F1-score, precision, and recall on different evaluation sets

While there is no common protein target or ligand between different subsets, it is still possible for the model to be biased for proteins similar to those in the training set. To demonstrate that the model is not biased for specific groups of protein. The AUC-PR values of the model on sampled protein targets are not correlated with the protein sequence lengths (Fig 3A). Likewise, the AUC-PR performances are also not correlated with the similarity between the sampled evaluation set proteins and the training set proteins (Fig 3B). Overall, the model did not have significantly different performances across protein families (Fig 3C). Nonetheless, the model showed the lowest AUC-PR on Peptidase S1 and TKL Ser/Thr protein kinase families, but the highest AUC-PR on Peptidase M1 and Tyr protein kinase families.

**Fig 3.**
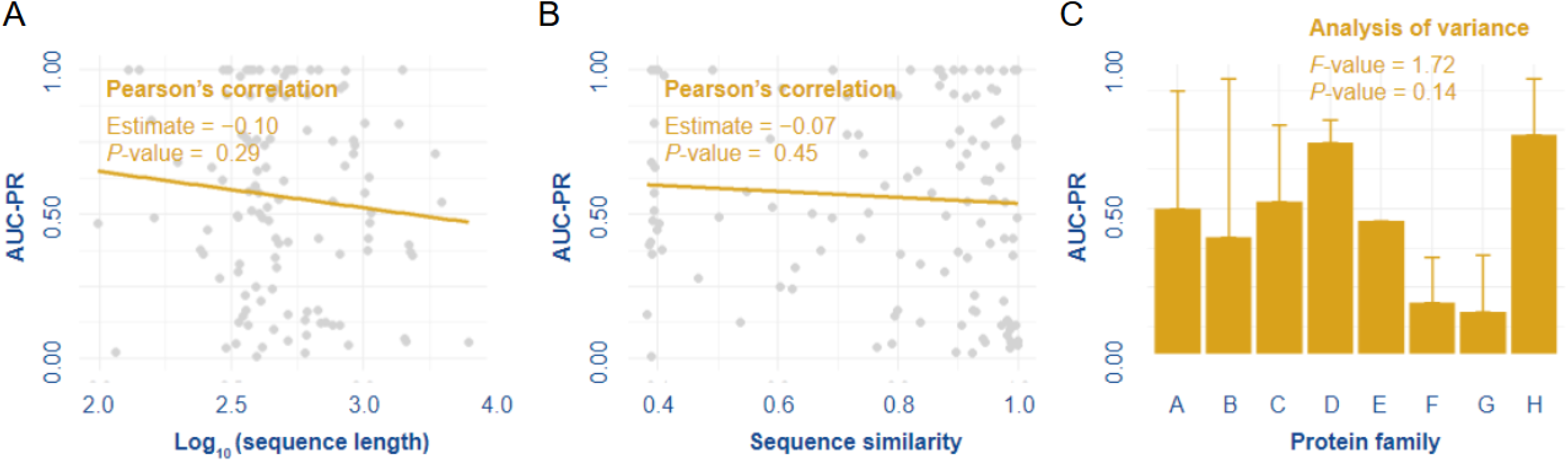
Model is generalizable to proteins unseen by the model. (A) There is no correlation between prediction performance (AUC-PR) and sequence length (log) of protein sequence. (B) There is no correlation between prediction performance and similarity of the predicted sequence relative to the sequence in the training set. (C) Prediction performance is not significantly different across protein families.

To demonstrate that prediction score or the probability of strong binding may also reflect the level of binding affinity, we map the true pIC_50_ values to the predicted scores of protein-compound pairs in the evaluation set. Interestingly, even if the model was strictly trained using a binary focal cross-entropy loss and a small range of continuous soft labels, the model was able to capture the continuous nature of binding affinity as indicated by a significant correlation coefficient of 0.4598.

### Virtual screening and molecular docking of ASII on NS1

The binding between NS1 of four DENV serotypes and a total of 177 natural compounds from Sigma-Aldrich were predicted. The results of top 10 compounds were given as examples in Table 2. Astragaloside II (ASII) was selected for the lack of report on severe adverse effects.

**Table 2.** Top 10 Sigma-Aldrich natural products pre-screened by our model.

| **PubChem substance ID** | **Synonym(s)** | **Type 1** | **Type 2** | **Type 3** | **Type 4** |
|---|---|---|---|---|---|
| 329824822 | Coloside A | 0.7057 | 0.7049 | 0.7062 | 0.7059 |
| 329824963 | Aconitone derivative | 0.6790 | 0.6784 | 0.6796 | 0.6791 |
| **329824948** | **Astragaloside II** | **0.6109** | **0.6099** | **0.6113** | **0.6111** |
| 329824750 | Tunicamycin A | 0.6091 | 0.6085 | 0.6095 | 0.6092 |
| 329824975 | Astramembrangenin | 0.6005 | 0.5995 | 0.6008 | 0.6007 |
| 24896458 | Dictamnolactone | 0.5805 | 0.5796 | 0.5813 | 0.5807 |
| 329824951 | Mesaconitine | 0.5677 | 0.5667 | 0.5679 | 0.5678 |
| 329824950 | Astragaloside I | 0.5605 | 0.5590 | 0.5606 | 0.5606 |
| 329824957 | Triptonide | 0.5558 | 0.5548 | 0.5565 | 0.5559 |
| 329824807 | Anacardic acid C15:3 | 0.5516 | 0.5505 | 0.5519 | 0.5518 |

Values are the model’s predicted probabilities of strong binding to NS1 of the four DENV serotypes (types 1–4). Astragaloside II (bold) was selected for follow-up.

CBDock2 [16] is one of the latest publicly available docking algorithms, which improves upon the existing AutoDock Vina 1.1.2. by first guiding the molecule via cavity detection based on both protein-surface-curvature and homologous template alignment. ASII was predicted by CBDock2 to bind to NS1 dimer (PDB: 4O6B [17]) with the two top affinities of -8.5 kcal/mol and -7.3 kcal/mol respectively. The binding sites of the two top affinities are located in the dimeric junction of the ventral side (the opposite side with respect to the β-roll) and in the crevice between the β-ladder end and the wing domain (Fig 4). Afterward, the identified pocket of the highest affinity was used for the validation with 3 experimentally verified controls [19] by SwissDock [18].

**Fig 4.**
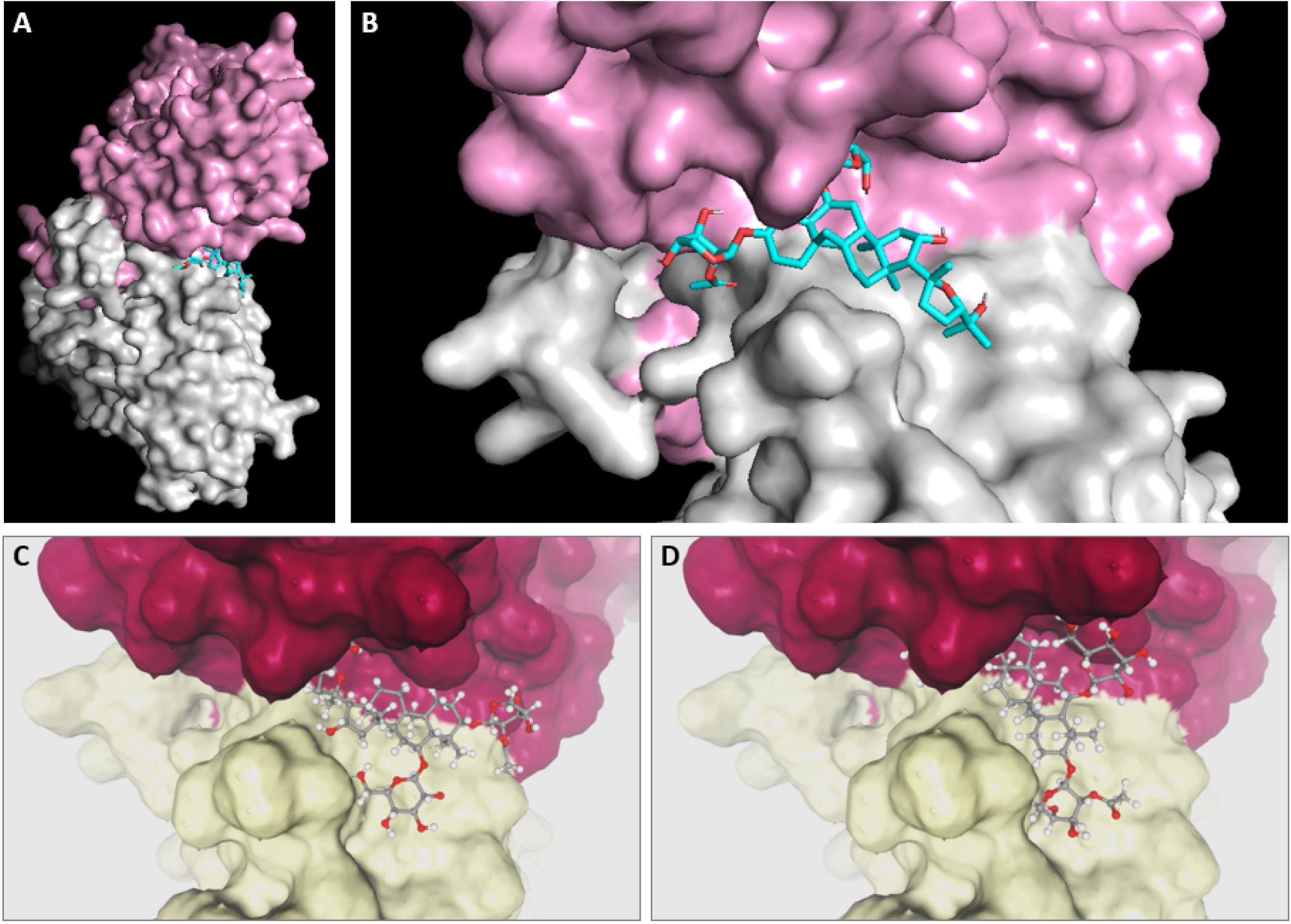
Molecular docking simulation of ASII on NS1 dimer. (A, B) Overall NS1-ASII complex predicted by CBDock2 with ASII binding site located in the dimeric junction on the ventral side, opposite to the β-roll. (C, D) The two highest affinity binding sites of ASII to NS1 (−8.0618 and -8.4975 respectively) predicted by SwissDock Attracting Cavities 2.0. given the binding pockets predicted by CBDock2. The two homo-subunits are highlighted in different colors (pink vs light gray; reddish pink vs pale yellow).

SwissDock-predicted binding affinities of the 3 controls match the order of the IC50 measured as inhibition of IL-6 release in PBMC activation assay. While SwissDock predicted ASII binding to be stronger than C01 but weaker than both C12 and C012, our model predicted ASII to be stronger than all the control ligands (Table 3).

**Table 3.** Comparison between the previous report, the model prediction and SwissDock binding scores of ligands on NS1 of DENV-2.

| **Ligand** | **pIC<sub>50</sub> for IL-6 induction** | **Model prediction** | **SwissDock score** |
|---|---|---|---|
| C012 (PG545) | 6.676 | 0.4892 | −11.8055 |
| C01 | N/A | 0.4600 | −7.6106 |
| C12 | 4.535 | 0.3997 | −9.7654 |
| ASII | – | 0.6098 | −8.0523 / −8.4975 |

pIC<sub>50</sub> values for inhibition of NS1-induced IL-6 release in PBMCs are from Modhiran et al. [19]. N/A, no inhibition observed.

### ASII inhibits NS1-induced endothelial hyperpermeability without cytotoxic effects

HMEC-1 monolayers were treated with varying concentrations of ASII to assess its impact on NS1-induced cell hyperpermeability and cytotoxicity (Fig 5). HMEC-1 cells were grown on Transwell permeable membranes as an *in vitro* model of barrier function and incubated with DENV-2 NS1 and/or ASII. Endothelial cell permeability was examined by measuring TEER at different time points over 24 hours, are shown in Fig 5A. DENV-2 NS1 alone induced endothelial hyperpermeability, significantly reducing the relative TEER to 0.56 ± 0.01 (mean ± standard error of the mean [SEM]) at 9 hours post-exposure, compared to control cells (*P* < 0.001). The relative TEER of NS1-exposed cells recovered to near baseline by 24 hours.

**Fig 5.**
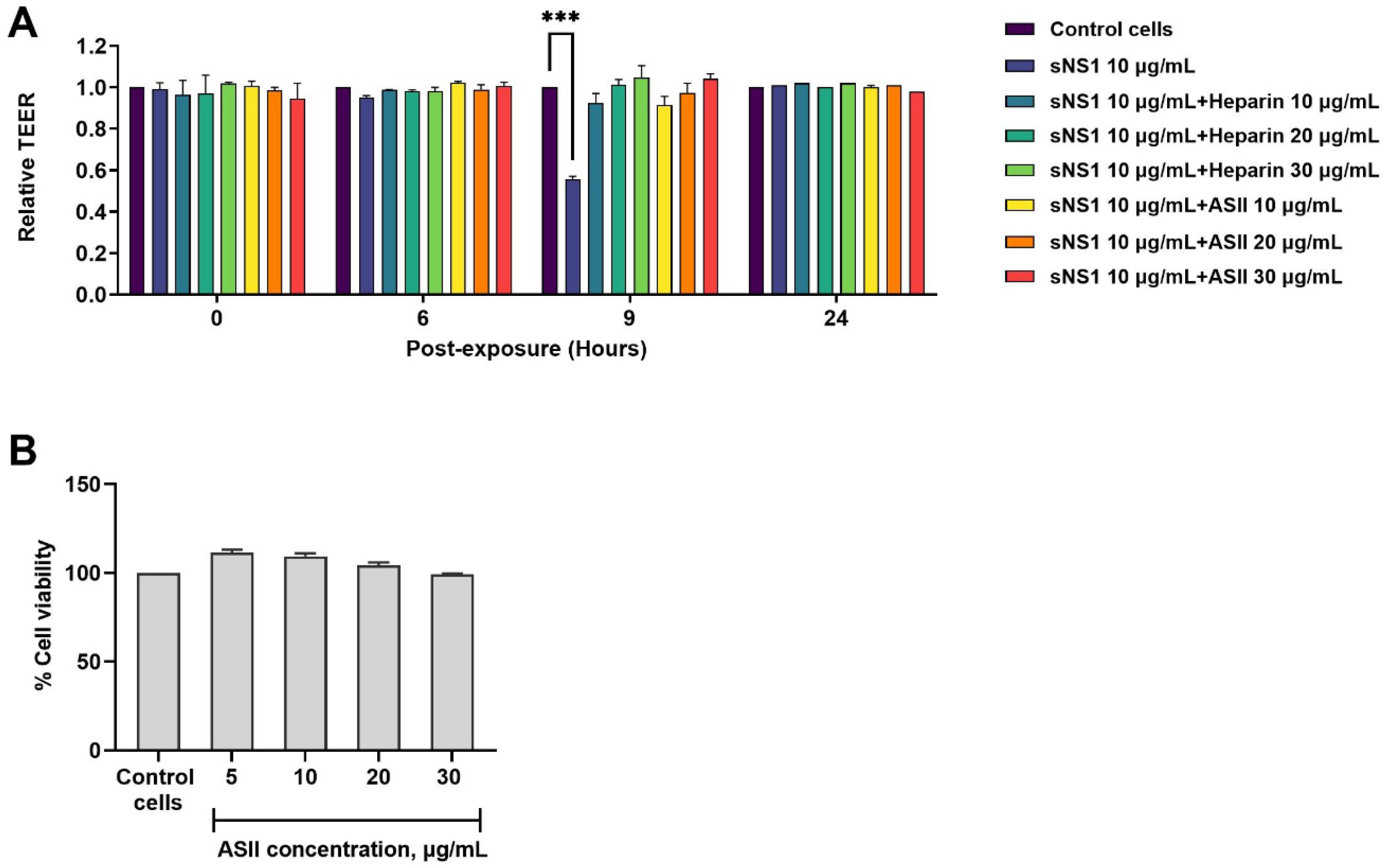
Inhibitory effect of ASII on NS1-induced endothelial hyperpermeability without evident cytotoxicity. (A) HMEC-1 monolayers were grown on 24-well Transwell inserts, and DENV-2 NS1 were added to the apical chamber to induce endothelial hyperpermeability. Relative TEER values were measured at the indicated time points over 24 hours for experiments evaluating the effect of ASII at concentrations of 10, 20, and 30 μg/mL on NS1-induced endothelial hyperpermeability, and experiments with heparin, used as a positive control. Statistically significant differences between experimental groups and the control group were determined using an ordinary two-way ANOVA with Dunnett’s multiple comparisons. Asterisks denote significance level: \*\*\**P* < 0.001. (B) The viability of HMEC-1 treated with ASII was measured to evaluate the cytotoxic effects of ASII at concentrations of 5, 10, 20, and 30 μg/mL. ASII did not exhibit cytotoxicity in HMEC-1, as the viability remained comparable between untreated control and ASII-treated cells. (A-B) Bar graphs with error bars represent the mean ± SEM from three independent experiments.

In contrast, when treated with ASII at concentrations of 10, 20, and 30 μg/mL, the mean relative TEER ± SEM across NS1-exposed HMEC-1 monolayers at 9 hours post-exposure were 0.92 ± 0.04, 0.97 ± 0.05, and 1.04 ± 0.02, respectively, which were not significantly different from those of control cells. This suggests that ASII mitigated endothelial hyperpermeability induced by DENV-2 NS1. Consistently, heparin, known to bind NS1, was used as a positive control, and comparisons of relative TEER values between control cells and NS1-exposed HMEC-1 treated with heparin at 10, 20, and 30 μg/mL revealed no significant differences at any time point. The TEER results for HMEC-1 treated with ASII or heparin alone are provided in S1 Fig, showing that neither ASII nor heparin alone affected HMEC-1, as the relative TEER remained similar to those of control cells.

To assess ASII cytotoxicity, ATP levels were measured to determine the viability of HMEC-1 treated with ASII. The cytotoxicity results for different ASII concentrations (5, 10, 20 and 30 μg/mL) treated on HMEC-1 are shown in Fig 5B. None of the tested concentrations of ASII exhibited significant cytotoxic effects on HMEC-1 at 24 hours post-treatment compared to untreated control cells.

### Effect of ASII on the accumulation of NS1 in HMEC-1

We investigated whether ASII reduces intracellular NS1 accumulation in HMEC-1. Cells were treated with NS1 and ASII, and intracellular NS1 levels were quantified using fluorescence intensity. At 9 hours post-exposure, the fluorescence intensities in NS1-exposed cells treated with ASII were not statistically different from those in cells exposed to equivalent amounts of NS1 without ASII treatment (Fig 6).

**Fig 6.**
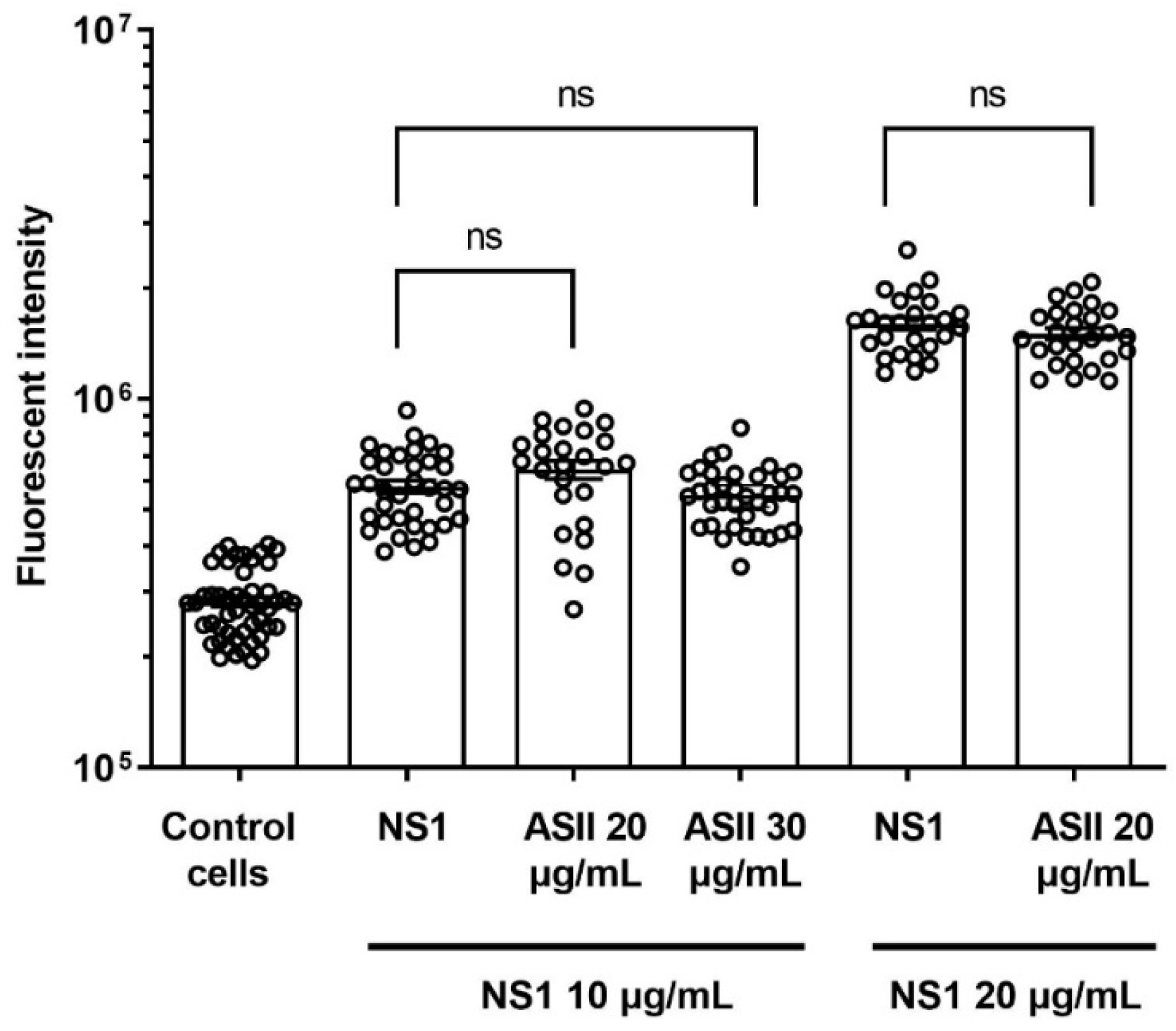
Internalization of DENV-2 NS1 in HMEC-1 treated with or without ASII. Bar graphs with error bars represent the mean ± SEM fluorescence intensity from two independent experiments. No significant differences in fluorescence intensity were observed between NS1-exposed cells treated with ASII and those without ASII treatment (*P* > 0.05), as determined by the unpaired *t*-test. ns; non-significant.

### Effect of ASII on NS1–HS binding

We assessed the ability of ASII to inhibit the interaction between NS1 and HS using a competitive ELISA, as NS1 has been shown to interact with HS on endothelial cell surfaces [8]. Specifically, NS1 (3 μg/mL) and varying concentrations of ASII (1 – 100 μg/mL) were added to wells coated with HS. As shown in Fig 7, ASII inhibited the binding of NS1 to HS in a dose-dependent manner, with an IC_50_ of 51.12 μg/mL (95% confidence interval [CI]: 47.91 – 56.24 μg/mL). These results indicate that ASII competes with NS1 for binding to immobilized HS, potentially by directly interacting with NS1. Based on the dose-response curve, interpolation from the best-fit model predicts that ASII at 10, 20, and 30 μg/mL inhibits NS1-HS binding by 7.96% (95% CI: 5.86–10.06%), 13.78% (95% CI: 11.61–15.95%), and 26.33% (95% CI: 23.58–29.07%), respectively.

**Fig 7.**
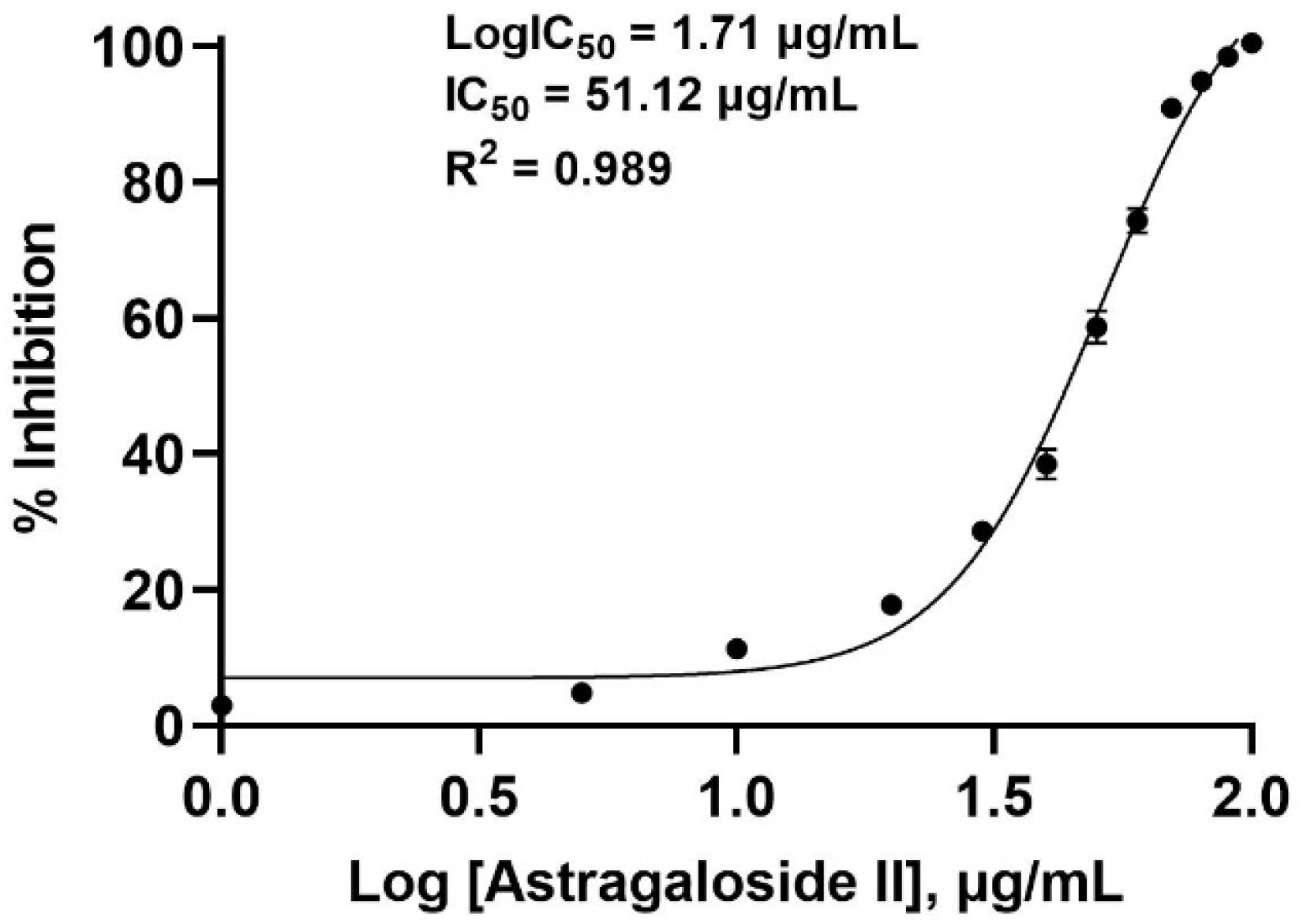
Dose-response curve and IC_50_ determination of ASII for inhibiting NS1-HS binding. The curve was generated using a nonlinear regression model. The line graph with error bars represents the mean ± SEM from two independent experiments.

### Potential effects of ASII on NS1–TLR4 binding

We hypothesized that ASII inhibits binding NS1-TLR4 binding and subsequently prevent TLR4 signaling-associated hyperpermeability. AlphaFold3, the state-of-the-art model to predict binding complex between protein and macromolecule of choices, was used to identify potential binding sites between the two proteins. AlphaFold3 [20] was able to predict NS1 dimeric structure with remarkable similarity to the PDB structure 4O6B used in molecular docking of both SwissDock and CB-Dock2, supporting its potential for the prediction of NS1-TLR4 binding sites (Fig 8). The two predicted binding sites were located on the junctions between each subunit. Interestingly, the dorsal TLR-4 binding sites, which are on the ventral side of NS1 as NS1 is positioned upside-down relative to what is shown in Fig 4, also overlap with the highest affinity binding sites of ASII predicted by CB-Dock2 and the potential sulfate binding sites [21,22].

**Fig 8.**
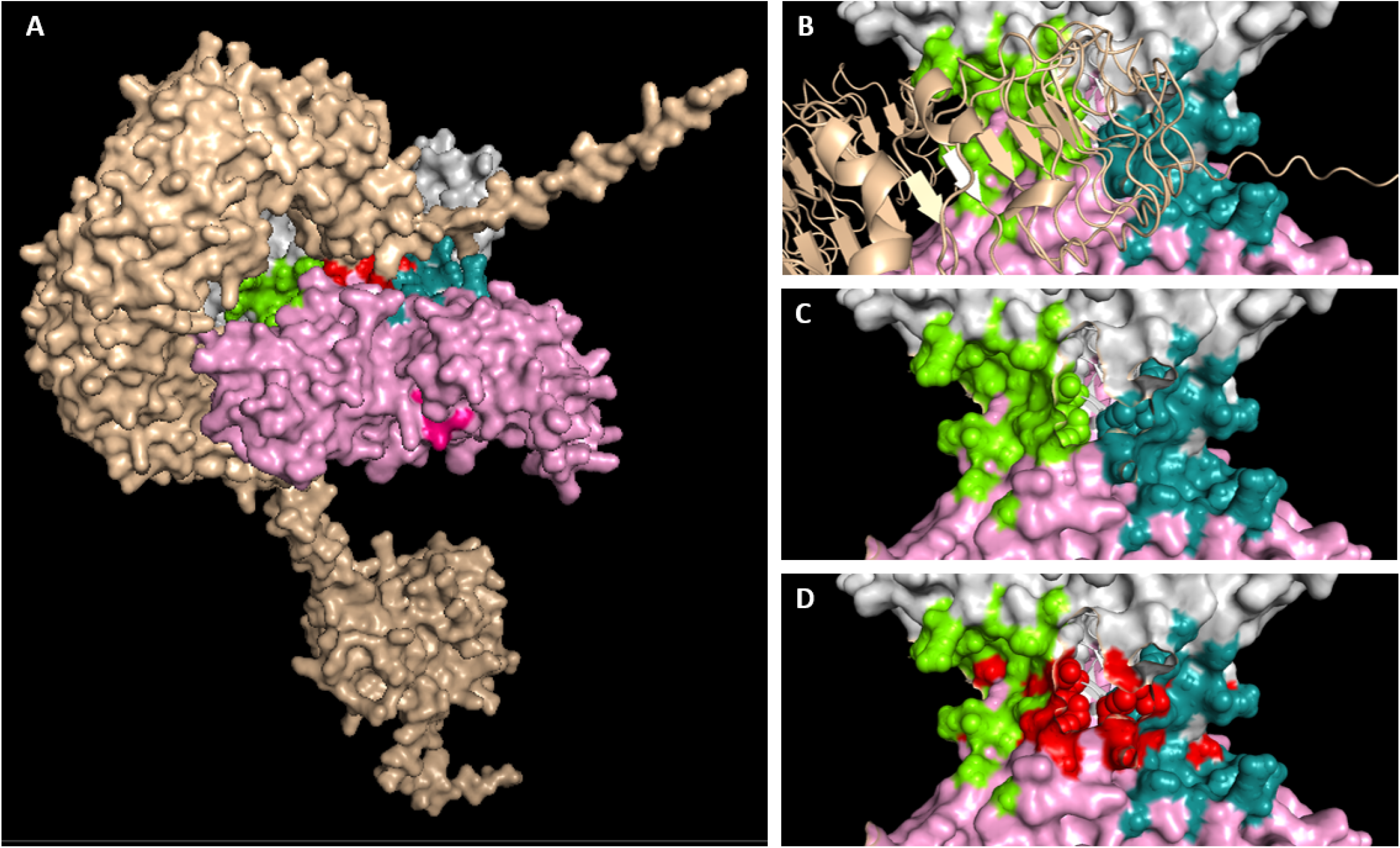
AlphaFold3-predicted 3D complex structure between NS1 dimer and TLR4 with potential ASII binding sites. (A) Overall structure of the complex with the two binding sites of ASII with the same highest affinity by CBDock2 (chartreuse green and deep teal), sulfate binding residues (red), GDI sequence of the wing domain (reddish pink). The two homo-subunits of NS1 are depicted in different colors of pink and light gray. TLR4 is in wheat yellow. (B, C, D) Top view of the structure to show the overlaps between TLR4-NS1 direct binding region (indicated by loss of surface render), NS1-ASII-binding regions, and sulfate-binding residues.

The two sites (chartreuse green and deep teal) are the same, but of C2 symmetry of the other. This potential inhibitory site also overlaps with the site predicted by SwissDock. The GDI sequence has been reported to confer endothelial cell binding specificity and vascular leakage [26]. Interestingly, the second highest affinity binding of -7.3 kcal/mol of ASII predicted by CBDock2 also involves D91 and I92 of the GDI motif.

## Discussion

The improved accuracy of our model compared with the original CNN-based DeepConvDTI can be attributed to several advantages. First, the stacked CNN architecture’s ability to learn hierarchical features from protein sequences and ligand fingerprints enables it to capture subtle patterns in binding interactions. Regularization by mish and Batch normalization helps reduce model overfitting especially for the training of large dataset. Lastly, using rectified Adam helps reduce variance of adaptable learning rate across the layers and stabilize the model further. Furthermore, one key factor that enhances model generalizability was the splitting algorithm that ensures no common protein or ligand was shared across different subsets. Although this splitting did not account for high but not exact similarity, the correlation analysis with respect to protein similarity within sample groups showed that the model did not preferentially perform better in evaluation set samples with higher protein similarity to the training set.

Among the top hits from our virtual screening, both Coloside A (prediction score: 0.7057) and an aconitane derivative (0.6791) showed higher predicted binding affinities than Astragaloside II (0.6108). However, safety considerations strongly influenced our lead selection. Coloside A belongs to the cucurbitacin family, which is associated with severe hepatotoxicity and gastrointestinal effects [27]. Similarly, the aconitane derivative represents a class of compounds known for their narrow therapeutic window and potentially lethal cardiotoxicity [28]. In contrast, Astragaloside II, a cycloartane-type triterpene glycoside from *Astragalus membranaceus*, has demonstrated a favorable safety profile through its long history of traditional medicine use [29].

NS1 is recognized as an attractive therapeutic target for modulating the endothelial barrier during dengue infection. In our search for compounds that target NS1, ASII—a saponin purified from Radix Astragali, the dried root of *Astragalus membranaceus*, used in traditional Chinese medicine—has emerged as a promising candidate. ASII has been reported to modulate the immune response and inhibit tumor growth [30]. Recent computational docking and *in vitro* studies suggest that ASII exhibits substantial antiviral activity against DENV [31]. Our compound-target interaction prediction model showed a high binding affinity between ASII and NS1, though how this interaction reduces endothelial hyperpermeability has not yet been investigated. In this study, we investigated whether ASII binds to NS1 and inhibits NS1-induced endothelial hyperpermeability. TEER measurements and immunofluorescence confocal microscopy revealed that ASII preserved the endothelial barrier in NS1-exposed HMEC-1 but did not prevent NS1 internalization. Competitive ELISA further showed that ASII competed with NS1 for binding to immobilized HS, with a high IC_50_. These findings suggest that ASII may inhibit NS1-induced hyperpermeability by disrupting downstream signaling pathways of NS1, rather than blocking its internalization.

Endothelial permeability is regulated by two key factors: 1) the EGL, which forms a protective membrane on the endothelial cell surface, and 2) adhesion and tight junctions, which maintain endothelial integrity. Since ASII preserved endothelial barrier without blocking NS1 internalization, its protective effect likely occurs downstream of internalization. We hypothesize that ASII does not interfere with NS1 uptake but instead modulates signaling pathways involved in endothelial dysfunction, such as TLR4-induced macrophage migration inhibitory factor (MIF) secretion, which increases active heparanase-1 levels [32], or NS1-mediated activation of matrix metalloproteinase-9 (MMP-9), which disrupts cell adhesion and tight junctions [5]. Alternatively, ASII may enhance junction stability or mitigate NS1-induced glycocalyx degradation, preserving TEER despite NS1 internalization.

Previous studies have demonstrated that NS1 can induce endothelial hyperpermeability through multiple mechanisms, including TLR4-mediated signaling [10], glycocalyx degradation [33], and clathrin-mediated endocytosis [34]. Modhiran et al. [10] reported that blocking TLR4 protected endothelial cells from NS1-induced hyperpermeability, suggesting that NS1-TLR4 interactions contribute to endothelial dysfunction. However, another study found that TLR4 activation alone is insufficient to cause hyperpermeability, emphasizing endothelial enzyme-mediated glycocalyx degradation as a key driver of NS1-induced vascular leakage [33]. More recently, it was shown that NS1 undergoes clathrin-mediated endocytosis, which is required for its effect on endothelial permeability, as blocking endocytosis prevented the reduction in TEER [34].

Structural prediction of the NS1-TLR4 complex by AlphaFold3 showed the overlap between NS1-TLR4 binding interface and predicted ASII binding sites and hence supported the notion that ASII binding may competitively prevent NS1 from binding to the TLR4. Furthermore, these binding sites overlap with the sulfate binding residues observed in the previous study [22]. Since SwissDock can successfully predict the binding affinity order of the sulfated glucose-rich C012 and C12 in the region, it is possible that ASII and C012 may inhibit NS1 with the same binding mechanisms as does heparan sulfate. Interestingly, ASII was also predicted to bind, albeit with lower affinity, to the GDI motif of the wing domain that was reportedly crucial to endothelial specific binding by NS1 [26]. However, GDI motif and this hypothetical ASII binding site are distant from the NS1-TLR4 binding interface. While GDI motif is required for NS1 binding to the endothelial cell, there was no evidence that it directly serves as active physical binding motif. The GDI may play a more important role in conformational mechanisms and flexibility of NS1 such that its binding to ligands may not affect its pathological functions.

This study has several limitations. Binding of ASII to NS1 was predicted computationally and inferred from a competitive ELISA; it was not measured with a direct biophysical assay. The ELISA and internalization experiments each rest on two independent experiments, and the TEER and viability data on three. Only one endothelial cell line, one NS1 serotype (DENV-2) and one commercially sourced compound were examined, and no infection or antiviral assay was performed, so the effect of ASII on DENV replication remains untested. The ASII concentrations that preserved TEER (10–30 µg/mL) inhibited NS1–HS binding by only 8–26% in the competitive ELISA, indicating that inhibition of HS binding alone is unlikely to explain the protective effect. The model also scored the inactive control C01 above the active control C12 (Table 3), so its ranking should be regarded as hypothesis-generating rather than confirmatory. Finally, TEER comparisons were made against untreated control cells; a direct comparison of NS1-exposed cells with and without ASII is needed to confirm protection.

In summary, this study indicates that ASII, which was predicted to bind NS1, preserved the endothelial barrier in NS1-exposed HMEC-1, although it did not prevent NS1 internalization. ASII may exert its protective effect on the endothelium by interfering with NS1-induced signaling and barrier disruption, rather than blocking NS1 uptake. Further studies are needed to determine whether ASII specifically inhibits pathways downstream of NS1 internalization, such as TLR4 activation, glycocalyx degradation, or cytoskeletal remodeling in endothelial cells.

## Supporting information

**S1 Fig.**
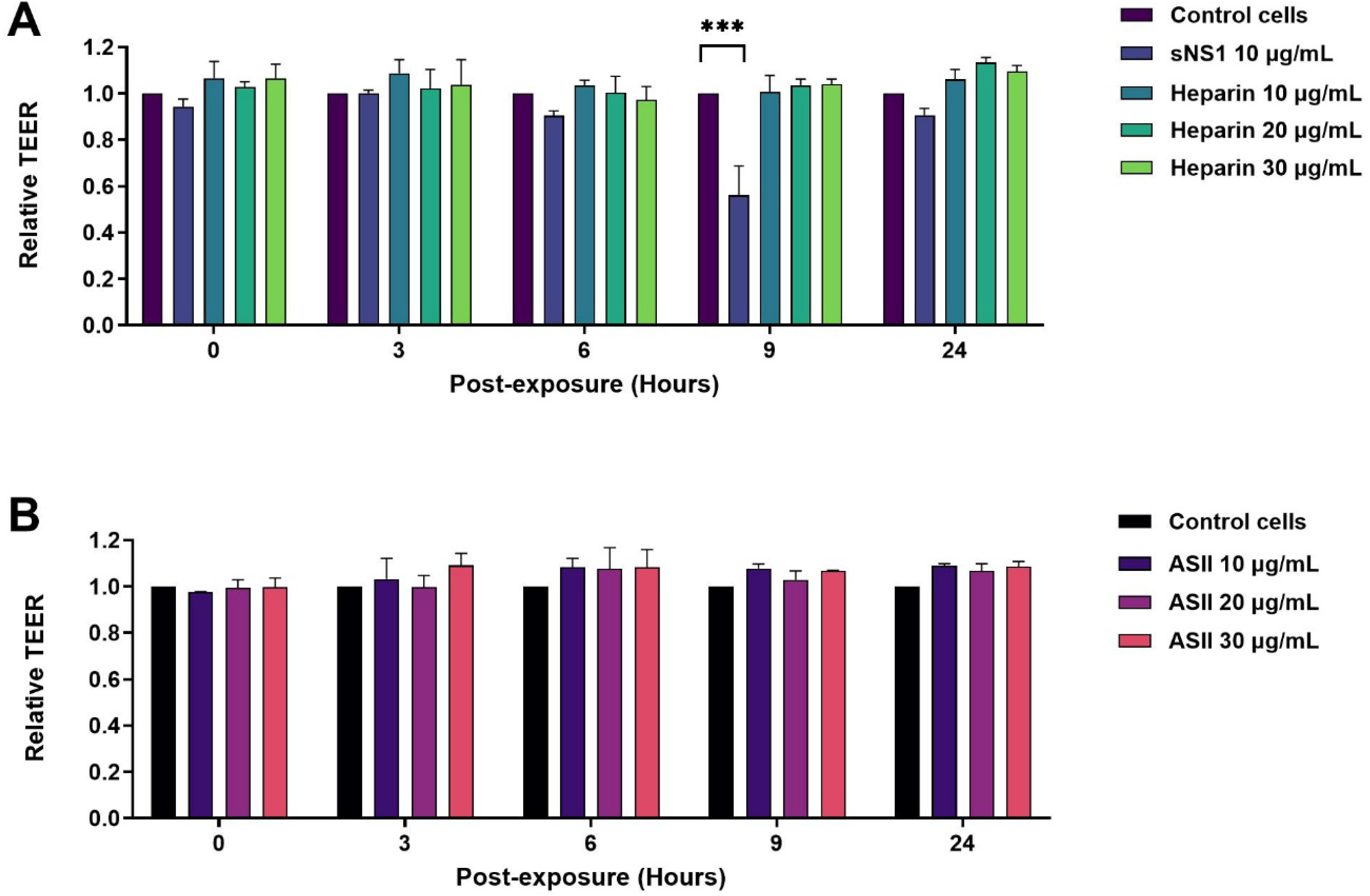
Relative TEER of HMEC-1 treated with heparin or ASII alone. (A) HMEC-1 monolayers treated with heparin (10, 20, or 30 µg/mL) and (B) HMEC-1 monolayers treated with ASII (10, 20, or 30 µg/mL) maintained relative TEER values that were not significantly different from those of control cells. Statistically significant differences between experimental groups and the control group were determined using an ordinary two-way ANOVA with Dunnett’s multiple comparisons. Asterisks denote significance level: \*\*\**p* < 0.001. Bar graphs with error bars represent the mean ± SEM from two independent experiments.

